# PhyloControl: a phylogeny visualisation platform for risk analysis in weed biological control

**DOI:** 10.1101/2025.06.11.658203

**Authors:** Stephanie H. Chen, Lauren Stevens, Ben Gooden, Michelle A. Rafter, Nunzio Knerr, Peter H. Thrall, Louise Ord, Alexander N. Schmidt-Lebuhn

## Abstract

1. Phylogenetic distance is a key measure used to develop species lists for host specificity tests that delimit the fundamental and realised host range of candidate weed biocontrol agents to meet the assumptions of the centrifugal phylogenetic method. Plant pathogens and insects, even those with broad host ranges, exhibit some degree of phylogenetic conservatism in their host plant associations. Thorough host-specificity testing is crucial to minimise the risk of off-target damage by biocontrol agents to native and economically important plant species. To facilitate this, host test lists need to be developed from an understanding of evolutionary relationships, usually visualised as a phylogenetic tree generated from genetic data, together with plant functional traits and geospatial information. Currently, the process of obtaining a host test list is not standardised, and the manual steps are time-consuming and challenging.
2. We introduce a user-friendly visualisation tool called PhyloControl to aid researchers in their decision-making during biocontrol risk analysis. PhyloControl integrates taxonomic data, molecular data, spatial data, and plant traits in an intuitive interactive interface, empowering biocontrol practitioners to summarise, visualise and analyse data efficiently.
3. Comprehensively sampled phylogenetic trees are often unavailable, and older published phylogenies often lack branch resolution and support, which increases uncertainty. PhyloControl includes a workflow implemented through Quarto notebooks in R that allows users to download publicly available DNA sequences and perform phylogenetic analyses. The modular workflow also incorporates species distribution modelling to predict the current and potential extent of target weed species.
4. PhyloControl will streamline the development of biocontrol host tests lists to support risk analysis and decision making in classical weed biological control.

## Introduction

In classical weed biological control, the release of a biocontrol agent is contingent on host-specificity testing to assess the risk of off-target damage in jurisdictions such as Australia, Canada, New Zealand, South Africa, and the United States of America. This risk assessment process seeks to ensure that agents are highly specific to the target weed. In accordance with the centrifugal phylogenetic method (Wapshere, 1974) and its modernisations (Briese, 2005; Sheppard et al., 2005), relatedness is an important factor in determining the host range of a candidate biocontrol agent. Closely related plants are more likely to have similar traits due to shared evolutionary history, and thus non-target plants closely related to the target weed are more likely to have similar or overlapping host recognition cues to candidate control agents (Jones et al., 2020, 2022; Le Falchier et al., 2025).

In the absence of formal analyses, estimates of relatedness may be based on ranks in the taxonomic classification (same genus, same tribe, same subfamily, etc.). Since growing numbers of phylogenetic analyses have been published, relatedness has often been expressed in degrees of phylogenetic separation, as the count of inferred lineage splits separating a target weed from its common ancestor with another species in the same phylogenetic tree (Kelch & McClay, 2004). An alternative metric of relatedness is patristic distance (also known as phylogenetic distance), which is the sum of branch lengths separating a target weed from another species across the phylogenetic tree. Patristic distance is preferred over degrees of separation as it can more accurately account for evolutionary relationships and variation in rates of evolution between species since the measure makes use of branch length information (Chen et al., 2024).

A host test list guides the specificity testing process and subsequent risk analysis as it helps define the scope of testing whilst prioritising species for testing (Chen et al., 2024). However, currently there is no standard approach to creating a host test list in weed biocontrol control, which can often be a difficult and long process, resulting in bottlenecks to the release of an agent to manage target weeds. Sequencing data and species occurrence records are becoming increasingly available in databases such as GenBank and the Global Biodiversity Information Facility (GBIF) and may be leveraged for biocontrol. DNA sequences can be used to understand evolutionary relationships between species through phylogenetic analyses, while occurrence data may be used for species distribution modelling and to visualise spatial overlap between species.

We introduce a modular workflow for generating inputs for visualisation via Quarto notebooks in R as well as an R Shiny application for visualisation. PhyloControl brings together genetic data in the form of a phylogenetic tree, spatial data, including species distribution modelling, and plant traits (Figure 1). The relatedness of species is summarised through phylogenetic distance measures, namely degrees of separation and patristic (phylogenetic) distance. This user-friendly decision support tool is designed for both biocontrol practitioners and regulators. We anticipate that PhyloControl will make the process of developing a host test list more efficient as well as transparent and reproducible.

**Figure 1.**
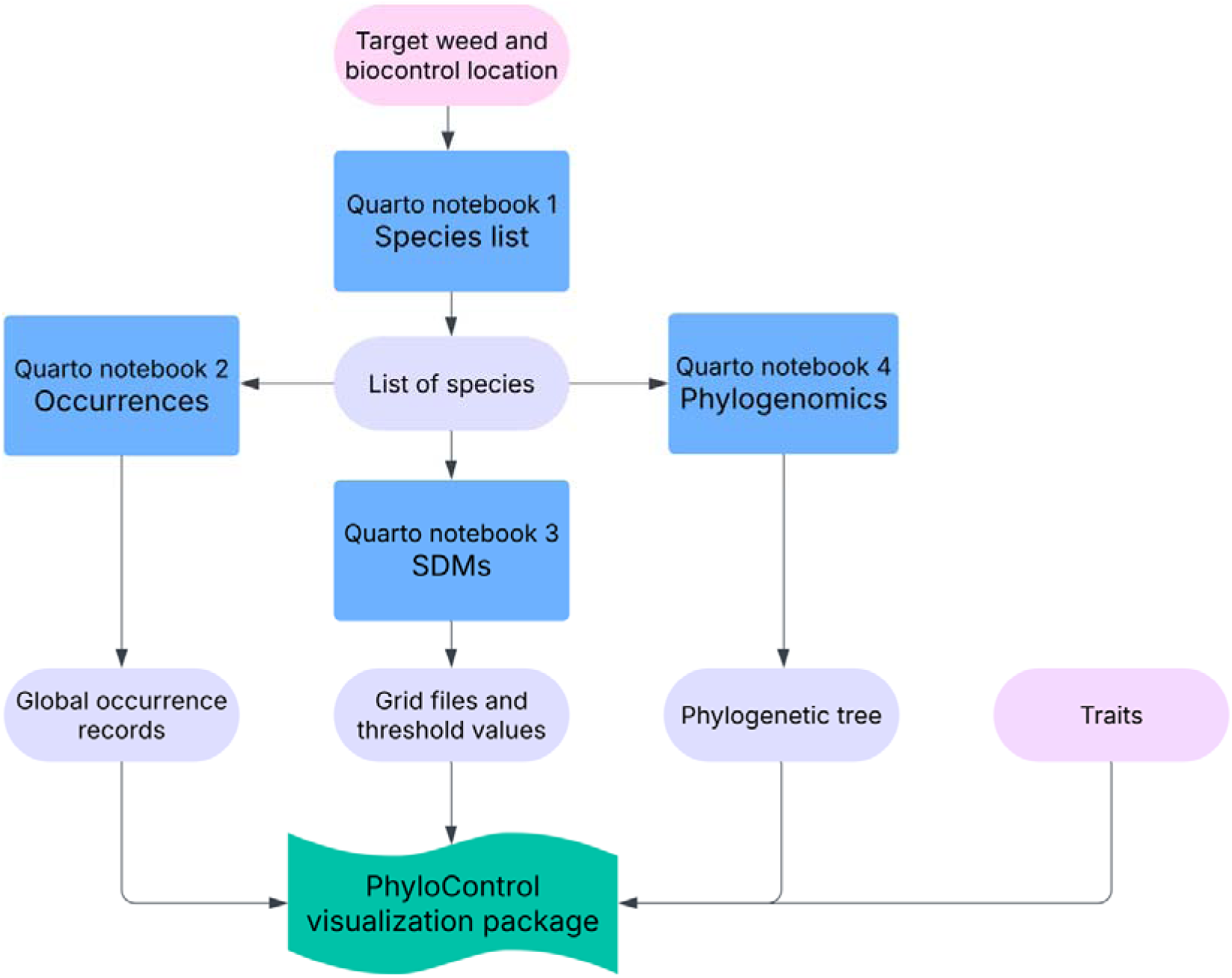
PhyloControl Quarto notebooks and visualisation app inputs. The scientific name of the target weed and country that the biocontrol project will take place in are needed as initial inputs for the first Quarto notebook to generate the species list. Flow diagram created in Lucid (lucid.co).

## Generating inputs for PhyloControl

A series of four Quarto notebooks has been developed to generate inputs for the visualisation app. The workflow is modular and reproducible which provides flexibility. For example, if a practitioner already has a species list or has generated data from their own sequencing experiment, these components can be skipped in the notebooks.

### Creating the species list (1_Species_list.qmd)

The kewr R package (Walker, 2024) is used to retrieve the scientific name of accepted species for a given plant family for specific location(s). The package uses Plants of the World Online, whose taxonomic backbone is the World Checklist of Vascular Plants. The output is a csv file of a list of species.

#### Downloading and filtering global point observation data (2_Occurrences.qmd)

Point observation data is retrieved from GBIF using the R package rgbif (Chamberlain & Boettiger, 2017). We chose to use preserved specimens (i.e., herbarium specimens), which have coordinates attached and are also expertly identified and curated. The coords2country function then attaches a country to each record for use in the R Shiny application.

### Species distribution modelling with MaxEnt and CLIMATCH (3_Species_distribution_modelling.qmd)

To predict the probability of occurrence for each species, we provide two methods for species distribution modelling: MaxEnt (Phillips et al., 2006), implementing the Java source code (Phillips et al., 2017), and CLIMATCH (Erickson et al., 2022). MaxEnt (maximum entropy modelling) is a widely used correlative machine-learning algorithm used for estimating habitat suitability based on presence-only data and environmental predictors. It is often chosen over other methods due to its predictive accuracy and ease of use (Merow et al., 2013). Meanwhile, CLIMATCH is a climate-matching tool that compares known species locations to locations to each possible invasion location and searches for the best match in a one-to-one association based on environmental variables, providing a conservative perspective compared to models that look for many-to-many associations. It requires the same presence-only data and environmental layers as input. It is a simpler approach compared to other SDM methods but is quick to run; government agencies in Australia and the United States use CLIMATCH as validation studies show that it is consistent predictor of invasion risk (Bomford et al., 2009; Hayes & Barry, 2008). The parameterisation of CLIMATCH intentionally resembles the CLIMATCH v2.0 webtool developed by the Australian Bureau of Agricultural and Resource Economics and Sciences (ABARES, 2020).

The climate layers used are the CHELSA v2.1 climatologies (Karger et al., 2017, 2021). Specifically, the variables starting with ‘bio’ that are not derived from other variables (i.e., bio1-19, excluding bio2-4) are accessed via the R package ClimDatDownloadR (Jentsch et al., 2023). These variables include mean annual air temperature (bio1), mean daily maximum air temperature of the warmest month (bio5), mean daily minimum air temperature of the coldest month (bio6), annual range of air temperature (bio7), mean daily mean air temperatures of the wettest quarter (bio8), mean daily mean air temperatures of the driest quarter (bio9), mean daily mean air temperatures of the warmest quarter (bio10), mean daily mean air temperatures of the coldest quarter (bio11), annual precipitation amount (bio12), precipitation amount of the wettest month (bio13), precipitation amount of the driest month (bio14), precipitation seasonality (bio15), mean monthly precipitation amount of the wettest quarter (bio16), mean monthly precipitation amount of the driest quarter (bio17), mean monthly precipitation amount of the warmest quarter (bio18), and mean monthly precipitation amount of the coldest quarter (bio19).

CLIMATCH is set to run for the extent of Australia by default, but may be used for other countries while MaxEnt is run at the global extent by default. For MaxEnt, the complementary log-log (cloglog) link function (Phillips et al., 2017) was used. Four threshold values were selected for visualisation – the maximum of Cohen’s Kappa (Monserud & Leemans, 1992), the maximum training sensitivity plus specificity (Cantor et al., 1999), threshold at which the sensitivity and specificity are equal, and sensitivity at 0.9. These were ordered and used as the boundary values for five bins.

The top two bins were coloured red on the map, the next was orange, and the next was yellow. The bottom bin remained transparent. CLIMATCH outputs a score between 0 to 10, indicating the degree of the match and results were visualised in a rainbow gradient palette.

### Building a phylogeny with public sequence data from GenBank 4_Sequences_and_phylogenomics.qmd)

The R package phruta (Román-Palacios, 2023) is used to find and download the markers available in GenBank and align the DNA sequences. The markers chosen can be set by a threshold where the default is to use markers that are found in at least 30 % of taxa in the species list provided. The IPS package is used for phylogenetic analysis with RAxML (Stamatakis, 2006). This includes a rapid bootstrap analysis and search for the best-scoring maximum likelihood tree as a quick way of generating a tree that is compatible with a local computer. Users can also generate their own sequences and phylogenies using alternative workflows (e.g. Chen et al., 2024), and the resulting Newick-formatted tree file may be used as input for visualisation in the PhyloControl R Shiny application.

### Example use case with comparison of different starting species lists

The code contains an example for a biocontrol project on the target weed flaxleaf fleabane, (*Erigeron bonariensis*) (CSIRO, 2023). We also compared three different scenarios where different levels of sequencing data and prior knowledge about the ingroup were available.

For Scenario 1, we generated a species list containing all Asteraceae occurring in Australia. In total, this species list had 1,425 species. An outgroup from the sister family Calyceraceae, *Boopis anthemoides*, was added for rooting. Of those species, 563 were not found and returned errors when GenBank’s taxonomic backbone was queried. There were three markers available for at least 30% of the 863 taxa in the cleaned species vector – *matK* (57%), ITS1 (32%), and ITS2 (30%), resulting in a concatenated dataset of sequence data for 328 species. The overall execution time for the full maximum likelihood analysis (100 bootstraps) was 5.1 h with 8 threads on a MacBook Pro with an M2 chip.

For Scenario 2, we focused on the tribe level and started with the species list from a target sequencing experiment that was curated by an Asteraceae botanist (i.e. from the experiment described for Scenario 3). Out of its 286 species, 102 gave errors when queries against GenBank’s taxonomic backbone, leaving 184 species to be input into the gene.sampling.retrieve function from the phruta package. There were 3 markers sampled in at least 30% of taxa: ITS1 (87%), ITS2 (85%), 5.8S rRNA (81%), resulting in a concatenated dataset covering 132 species. The overall execution time for the full maximum likelihood analysis was 0.36 h.

For Scenario 3, we aimed to generate a species tree at the tribe level of the target weed, i.e., Asteraceae tribe Astereae, using the Angiosperms353 target capture bait kit which amplifies 353 nuclear genes. In this paper, we focus on the concatenated analysis, rather than the coalescent analyses. This approach of using target capture sequencing allowed for comprehensive sampling and high phylogenetic resolution. The final tree contained 279 taxa (some samples failed to sequence or were quality filtered out), including 4 outgroups from other tribes of Asteraceae. These analyses required High Performance Computing (HPC), and the resources required were detailed in the paper where the phylogeny was initially published (Chen et al., 2024).

## PhyloControl visualisation tool

The visualisation component of PhyloControl is an R Shiny application (Figure 2). The required inputs for visualisation are the phylogenetic tree with optional components being the occurrence data, species distribution models, and plant traits (Table 1). Instructional text is provided through the ‘Help’ button via rintrojs (Ganz, 2016).

**Figure 2.**
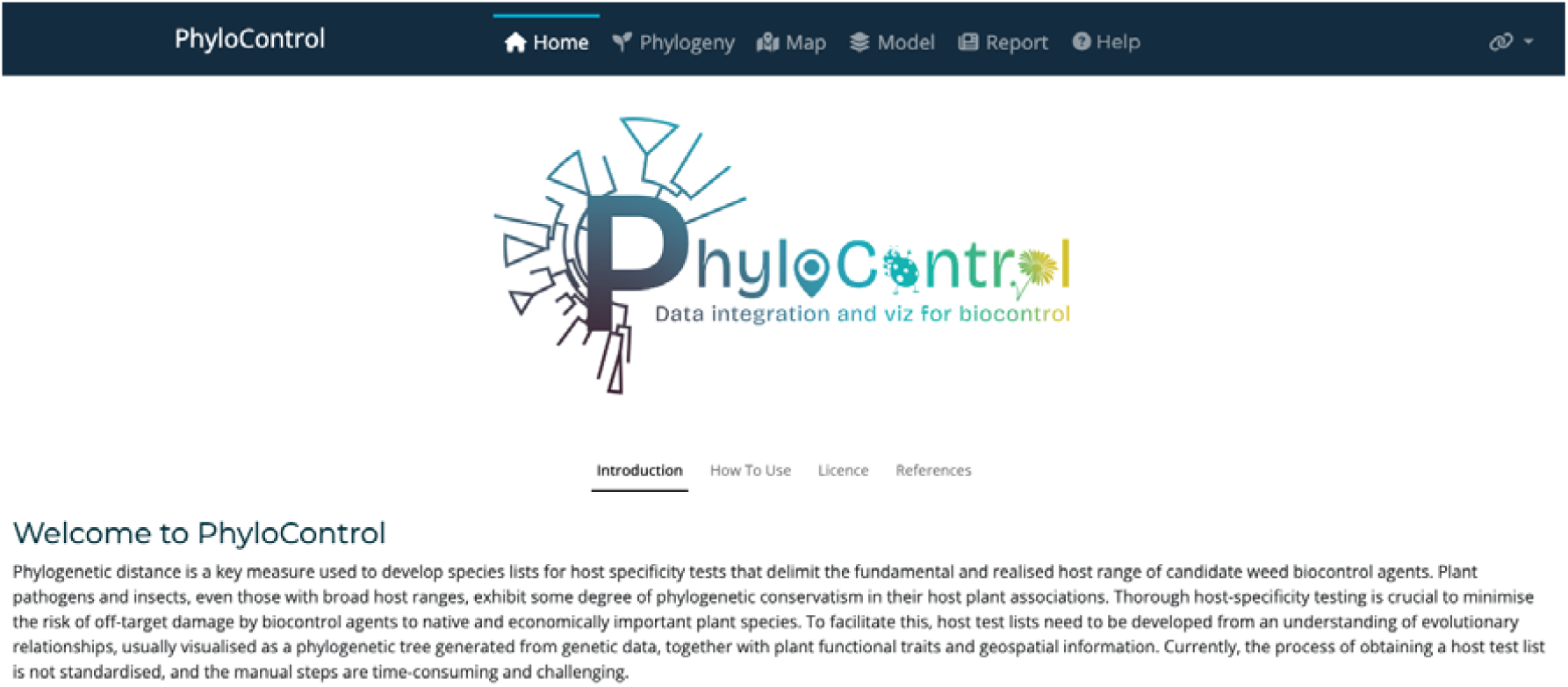
The home landing tab of the PhyloControl R Shiny application.

**Table 1.**
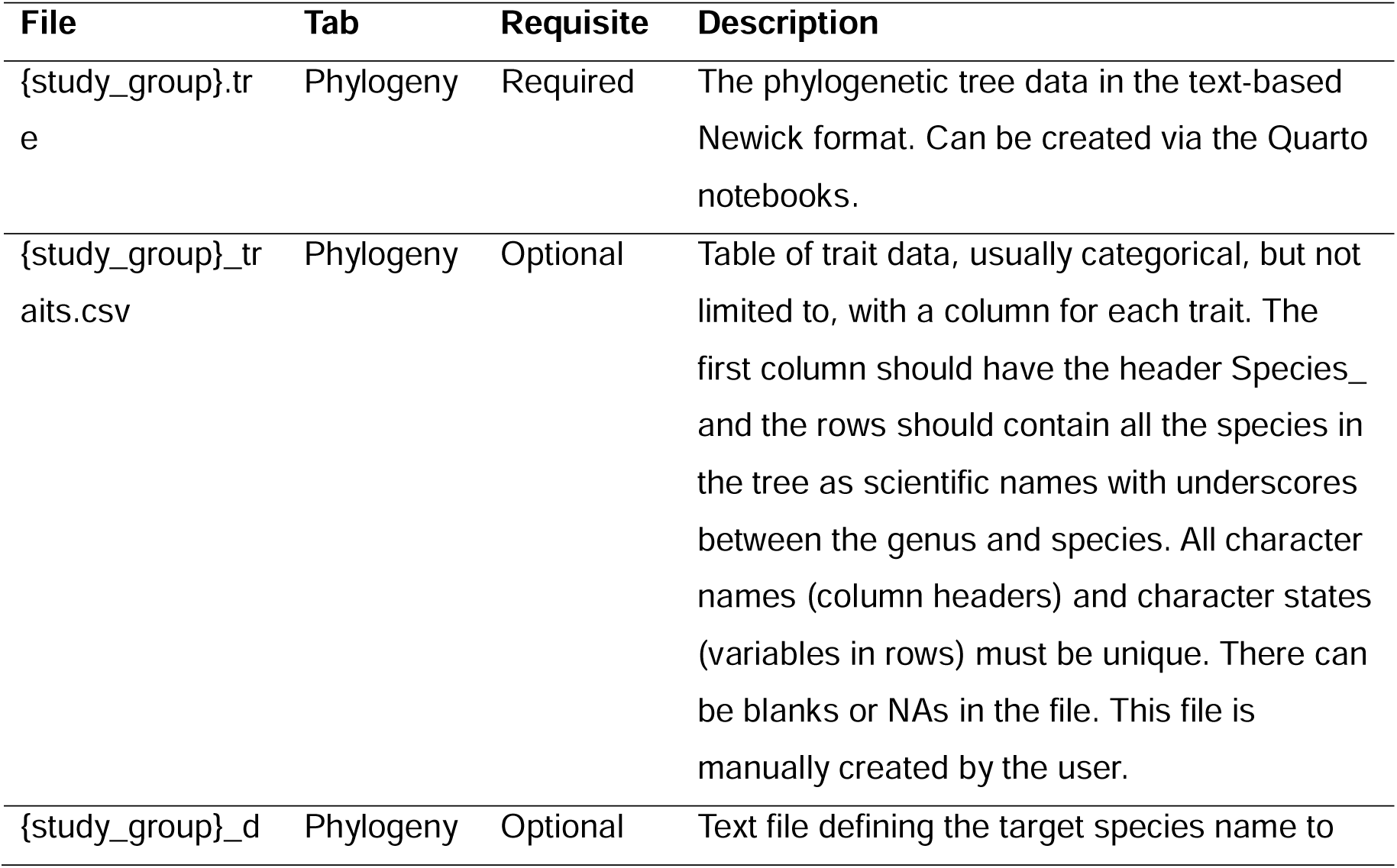
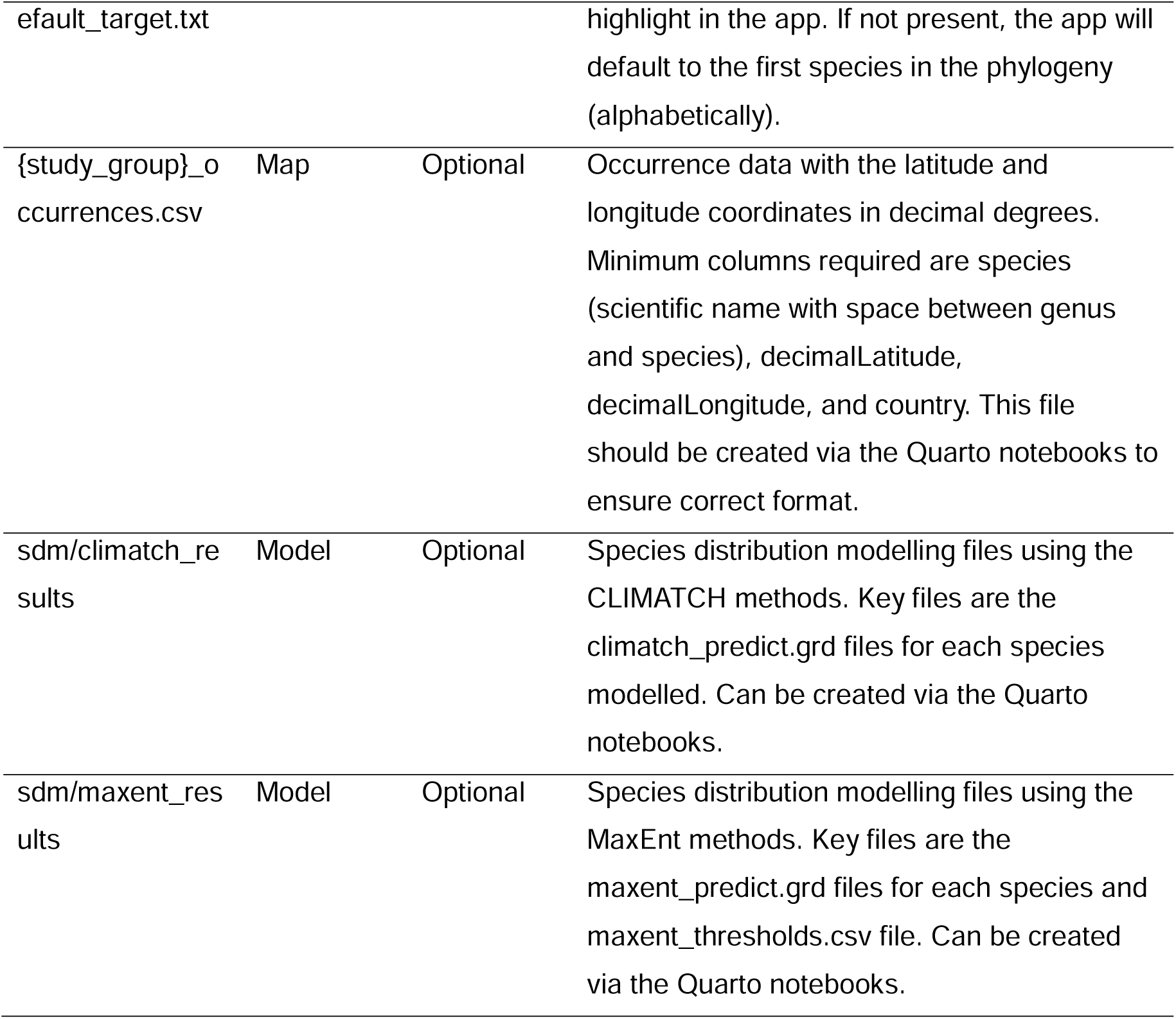
Inputs for visualisation with the PhyloControl R Shiny application.

The data displayed in the demo version of PhyloControl corresponds to Scenario 3 described above and is a previously published tribe level Astereae phylogeny using target sequence capture data (Chen et al., 2024). For the GBIF records, the initial query for preserved specimens with coordinate information contained 4,170,267 occurrences (GBIF.org, 2024). Following attaching country names and filtering, 85,679 records remained. For the species distribution modelling, the demo version of the app contains only the MaxEnt and CLIMATCH results for the target species. For demo data available on the DAP Portal, this is expanded to the six *Erigeron* species in the phylogeny. The modelling function took approximately a minute for each species. MaxEnt was run at a global scale while CLIMATCH was run only for Australia. A csv file with the native vs introduced status for each species in Australia was prepared manually.

### Phylogeny tab

The APE (Paradis et al., 2004) package was used for handling and manipulating the tree data while ggplot2 (Wickham, 2009), plotly (Sievert, 2020), and ggtree (Yu et al., 2017) were used for visualisation. There are options provided for the phylogeny for re-rooting, collapsing and expanding sections of the tree, and converting it to a cladogram. The phylogeny should be outgroup rooted using the most evolutionarily distant species for correct downstream inferences. Additionally, branch support values can be visualised as shades of grey. Where multiple support values are found in a Newick file, the visualisation will show the first set of values.

Up to four heatmaps as vertical bars may be simultaneously displayed to the right of the phylogenetic tree (Figure 3). Categorical and/or numerical traits provided via a csv file as well as the phylogenetic distance measures degrees of separation and patristic distance, which are calculated in app, may be shown. For categorical data, the discrete colour scale is limited to five categories to facilitate visualisation. The scico package (Crameri, 2018) was used for continuous scales. There is an option to highlight trait data in two colours: one colour for the traits with the same value as the target species and grey for all others. The visualisation may be downloaded as an image for use in reports, proposals, or manuscripts.

**Figure 3.**
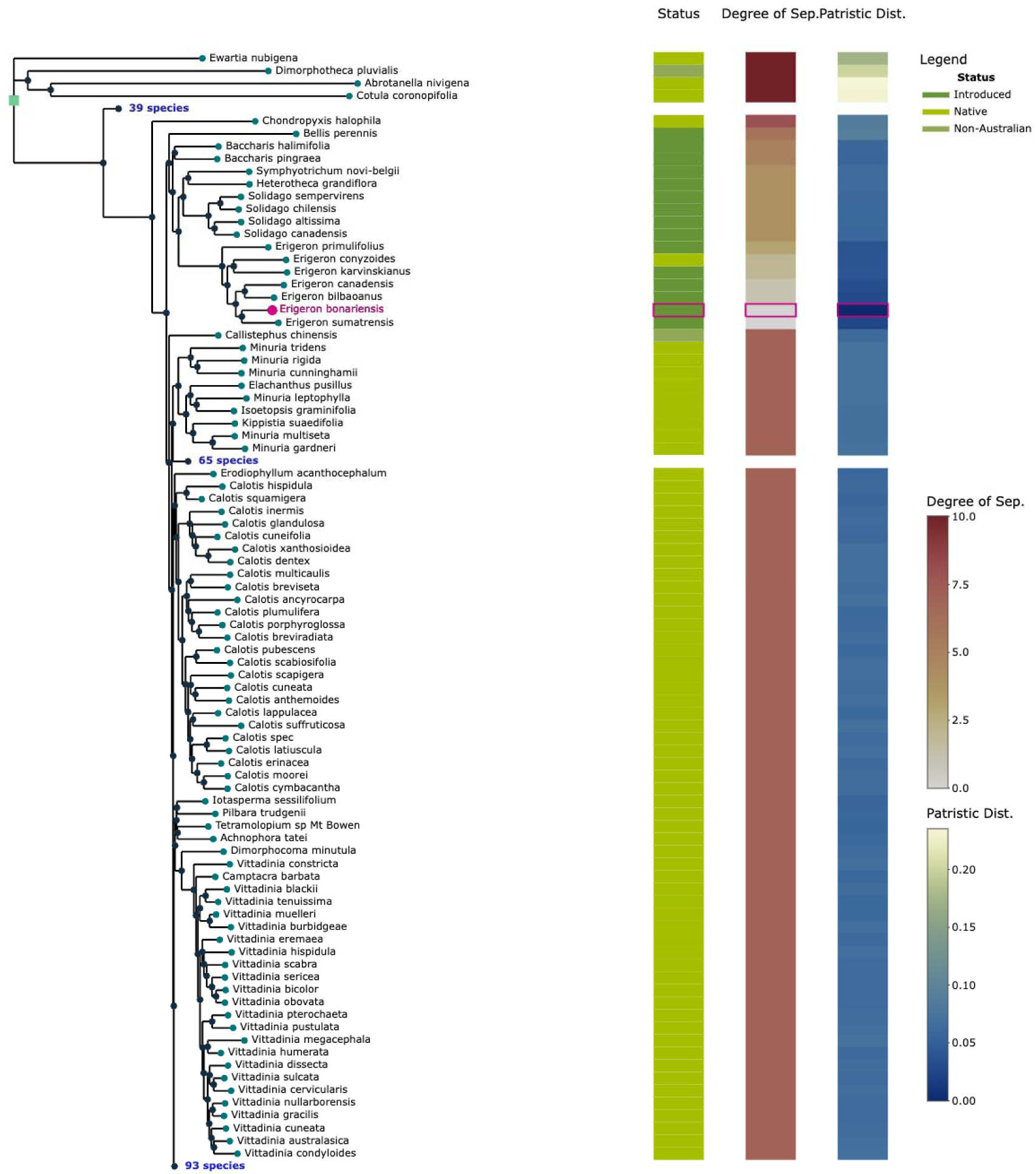
Phylogeny of the tribe Astereae for a biological control project on the target weed *Erigeron bonariensis*. Heatmaps display native vs introduced status in Australia, degrees of separation, and patristic distance. Note the clades that include all *Celmisia* (39 species), *Brachyscome* (65 species), and *Olearia* (93 species) were collapsed to shorten the length of the image.

### Map tab

Two species at a time can be selected to display occurrence records to assist in the visualisation of spatial overlap (Figure 4). A multiselect menu is available to filter the records to selected countries.

**Figure 4.**
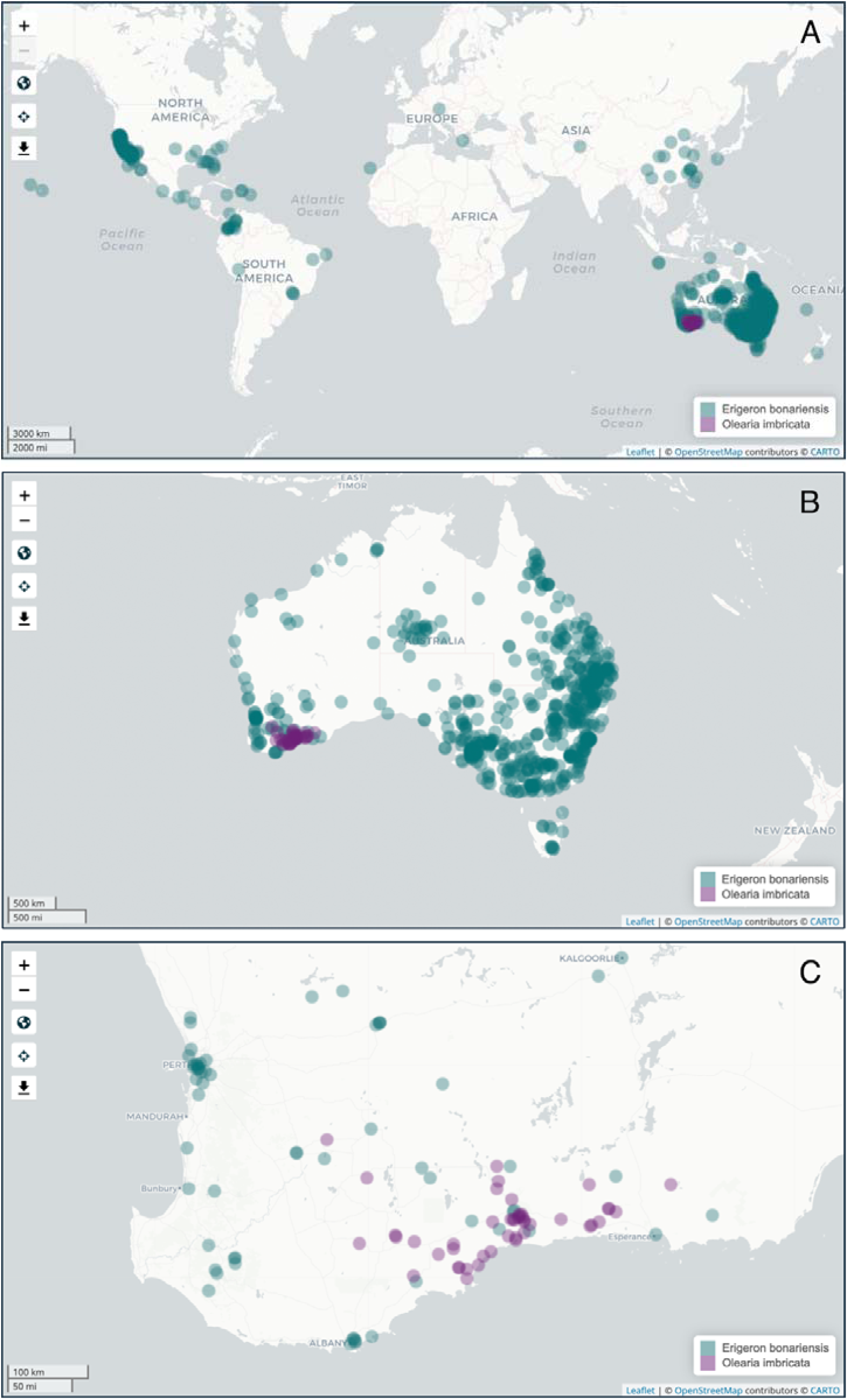
Global occurrence records from GBIF for the target weed flaxleaf fleabane (*Erigeron bonariensis*) in teal and the Australian native imbricate daisy bush (*Olearia imbricata*) in purple: A) All countries selected, B) Only Australia selected and C) Only occurrences in Australia displayed and zoomed to south-west Western Australia where the native daisy is endemic.

### Model tab

The results of species distribution modelling are displayed on a map coloured by MaxEnt presence of probability or CLIMATCH score, depending on the algorithm(s) chosen in the Quarto notebooks (Figure 5).

**Figure 5.**
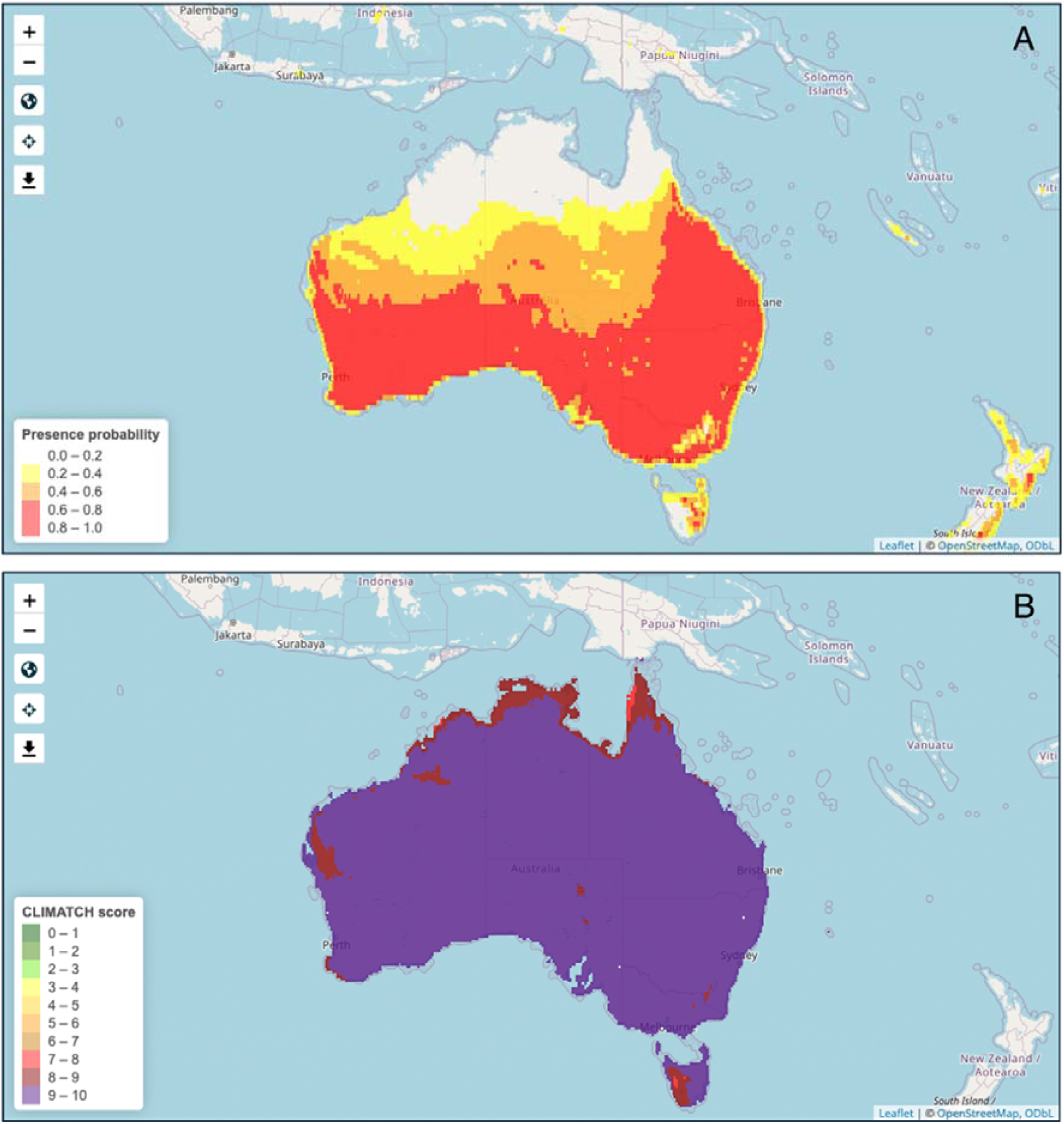
Species distribution model using A) MaxEnt and B) CLIMATCH for *Erigeron bonariensis* cropped to Australia.

### Report tab

A table of species in the tree ranked by the phylogenetic distance measures degrees of separation and patristic distance is available to download as a csv to assist a biocontrol practitioner to create a host test list.

## Discussion

PhyloControl is applicable to biological control projects globally, providing a reproducible and transparent means to bring together and visualise genetic, spatial, and trait data to optimise risk analysis. By focusing on species most closely related to the target weed, PhyloControl enables more targeted testing, reducing unnecessary experiments while allowing for thorough testing downstream in the biocontrol pipeline. Typically, a practitioner would be able to run the entire workflow and visualisation app on a local computer within a day, reducing the time and effort typically required to produce a host test list. This not only streamlines regulatory approval processes but also strengthens the scientific basis for biocontrol decisions which ultimately supports safer and more effective weed management strategies.

Gathering large volumes of data from disparate sources remains a challenge. Obtaining integrated taxonomic information is difficult and is a barrier to interoperability (Sandall et al., 2023). For example, the taxonomic backbones of the databases used for the Quarto notebooks, World Checklist of Vascular Plants, GBIF, and GenBank, do not align completely. Nonetheless, the R packages wrap their respective APIs to query and retrieve data facilitates the PhyloControl workflow. For the final trees for Scenario 1 and Scenario 2 of the *Erigeron* data, *Conyza*, a synonym of *Erigeron*, appeared in the tree, so the final trees may still require some manual curation if the study group has a recent history of taxonomic change.

A caveat of using publicly available sequencing data is that rare native species will likely be unsequenced, and these are some of the species important for biocontrol risk analysis where off-target damage to natives is a concern. As more genomics resources are generated, more comprehensive phylogenies will be able to be generated from GenBank data. In practice, biocontrol researchers will include threatened native relatives in the test list even if molecular data to understand evolutionary relationships are lacking. Here, PhyloControl allows for the identification of uncertain phylogenetic context so that important species may be targeted for additional sequencing.

Data availability also varies greatly across plant families. For example, families such as Fabaceae have had significant sequencing efforts directed towards them due to their economic importance (Group et al., 2013), so using GenBank data will result in a relatively comprehensive phylogeny. Checking for available GenBank data using the workflow may also help in the planning of a sequencing experiment to fill gaps and is a cost-effective way to obtain a comprehensive phylogeny. In general, genomics is being increasingly used in biological control in host testing lists and applications such as improving effectiveness of agents by targeting weed traits (Leung et al., 2020). Likewise, collecting trait data from open sources across the internet remains a challenge. Databases such as TRY (Kattge et al., 2020) and AusTraits (Falster et al., 2021) exist, but information will likely be patchy for a study group that a practitioner is interested in, and often they do not have global coverage, so this step remains manual in this initial version of PhyloControl. Currently, the user needs to identify informative traits that underpin agent-plant interactions and populated a spreadsheet with information with information from the literature.

Possible extensions of PhyloControl may include running the species distribution models with different climate change scenarios to understand effect of climate on invasion dynamics (Booth, 2018). Accessibility may be enhanced through the visualisation app being hosted through a server online if there is demand from practitioners. Additionally, whilst the tool is currently tailored to weed biological control, it may be modified to source information from other databases that are focused on the biological control of organisms other than plants.

PhyloControl addresses the challenges of a process that is currently unstandardised and time consuming. By integrating taxonomic, molecular, spatial, and trait data on target weeds and related plant species, PhyloControl empowers biocontrol practitioners to develop host test lists more efficiently, increasing confidence in risk analysis. Additionally, this improved approach has the potential to streamline regulatory by enhancing transparency and reproducibility. Therefore, we anticipate that the adoption of PhyloControl will lead to more robust and defensible risk assessments in weed biological control.

## Acknowledgements

We thank the attendees of the Annual Biological Control Workshop on 18 October 2023 at the Australian Government Department of Agriculture, Fisheries and Forestry for feedback on a prototype. The development of the R Shiny application was supported by the CSIRO Scientific Computing Collaboration Project programme.

## Data availability

The open source code for the PhyloControl R Shiny application is available at GitHub (https://github.com/csiro/phylocontrol-viz). The Quarto notebooks for generating the visualisation inputs are available at GitHub (https://github.com/csiro/phylocontrol-geninput). The outputs generated using the Quarto notebooks including the *Erigeron* demo dataset included with the app are available on the CSIRO Data Access Portal (https://doi.org/10.25919/21fr-hk78). A demo version of the app is available at https://shiny.csiro.au/phylocontrol-viz-demo/.

## Author contributions

Alexander Schmidt-Lebuhn, Ben Gooden, and Michelle Rafter conceptualised the project. Stephanie Chen and Nunzio Knerr prepared and tested the Quarto notebooks. The base framework concept and code were originally developed by Louise Ord with conversion to bslib done by Lauren Stevens. The core PhyloControl application code (Phylogeny and Map tabs) was written and implemented by Lauren Stevens based on the approach and design conceptualised by Louise Ord. Advice on solutions and implementation strategies for technical challenges were provided by Louise Ord and Lauren Stevens. The Model tab was created and developed by Nunzio Knerr and Stephanie Chen. The Report tab was created by Stephanie Chen. Alexander Schmidt-Lebuhn wrote the original functions for calculating degrees of separation and patristic distance. The *Erigeron* datasets used in this paper were prepared by Stephanie Chen, Nunzio Knerr, and Alexander Schmidt-Lebuhn. Stephanie Chen led the writing of the manuscript. All authors contributed critically to the drafts and gave final approval for publication.

## Conflict of Interest statement

The authors declare no conflict of interest.

## Notes

https://shiny.csiro.au/phylocontrol-viz-demo/

### Competing Interest Statement

The authors have declared no competing interest.

